# Mechanotranscriptomic Profiling of Breast Cancer Cells Intravasated from Engineered Microtumors

**DOI:** 10.64898/2026.02.28.708725

**Authors:** René Krüger, Miguel Fuentes-Chandía, Helly Atiya, Angeles de la Cruz García, Sadaf Pashapour, Aldo R. Boccaccini, Christine Selhuber-Unkel, Melanie Kappelmann-Fenzl, Anja K. Bosserhoff, Nicolás Tobar, Aldo Leal-Egaña

## Abstract

Intravasation is the process by which cancer cells breach the physical boundaries of a primary tumor and enter blood or lymphatic vessels. In this work, MCF-7 breast cancer cells were cultured within polymer-based microcapsules (here referred to as artificial microtumors) to investigate the transcriptomic and morpho-mechanical changes occurring in cancer cells during their release from these matrices, mimicking in vitro the process of intravasation.

Our results show that even confined and released cancer cells share approximately 95% of their global transcriptomic profiles, intravasation-like cells exhibited marked differences in the expression of pathogenic hallmarks, including pathways associated with cell proliferation, immunosurveillance, and dormancy. Notably, a clear upregulation of YAP/TAZ signaling was observed in released cells, a result further supported by single-cell traction force microscopy assays, demonstrating that those cells exhibit higher biomechanical activity compared to cells located within artificial microtumors or those cultured on conventional 2D flasks, as shown for intravasated cells in vivo.

To further enrich our investigation, the mechanotranscriptomic activity of MCF-7 cells was compared with suspended spheroids cultured on non-adherent surfaces (i.e., agarose hydrogels). Our results show that released cells displayed increased biomechanical activity and elevated expression of malignant markers, indicating that mechanical stress, beyond cell-cell contact alone, is required to trigger malignant responses. These observations were further supported by co-culture studies of MCF-7 cells with human fibroblasts and endothelial cells, which showed reduced proliferative and invasive capacities under confinement.

Overall, our findings demonstrate that shifts in mechanical and metabolic stress, as experienced during intravasation, act as critical stimuli driving mechanotranscriptomic programs associated with cancer progression.

## 1. Introduction

Among the different stages of cancer progression, intravasation refers to the process by which cancer cells breach the physical boundaries of primary tumors and enter blood or lymphatic vessels, thereby initiating the metastatic cascade.(^1–3^) This process involves remodeling of the tumor niche, driven by the secretion of matrix metalloproteinases and the exertion of forces required to mechanically disrupt the crosslinked protein matrix surrounding the cells.(^4–6^)

During intravasation, cancer cells undergo major morpho-mechanical changes, including extensive nuclear and cytoskeletal rearrangements, leading to chromatin condensation and nuclear compaction.(^7^) These processes are required to generate indentation forces capable of carving paths through the tumor stroma, with the nucleus acting as a piston.(^8–13^) As a consequence of the shear stress generated during cancer cell migration within a confined, semi-degradable matrix, the nuclear envelope frequently ruptures, becoming enriched in Lamin A/C and Lamin B1/B2.(^10–12^) Additionally, chromatin unravels, being exposed to shear and compressive stresses -typically experienced during cell migration and proliferation-, rendering DNA damage and mutations, and ultimately giving rise to heterogeneous cancer cell populations.(^12^)

Despite the substantial amount of research already conducted in this field, significant questions related to the intravasation process remain unanswered. Among these are the regulation and tuning of specific genetic and biomechanical malignant markers. Unsurprisingly, such information is difficult to assess in vivo, mostly due to the challenges of visualizing these processes in real time.

To address these limitations, several technical alternatives have been explored. Among them, the use of microfluidic systems seems to be the most widely used approach.(^14–16^) However, despite their broad adoption, these devices present notable drawbacks, including the need for highly trained personnel (restricting their implementation in public hospitals), limited scalability and adaptability during experiments, and the impossibility of implantation in small animal models, to enable the in vivo validation. In this context, the use of miniaturized tumors made of biocompatible polymers emerges as a promising strategy to overcome these technical challenges, offering a novel in vitro platform to study intravasation. We, among others, have extensively worked on the design and fabrication of polymer-based microtumors composed of alginate and gelatin, aimed to resemble the biophysical properties experienced by cancer cells when confined within primary tumors.(^17–22^)

Among the advantages offered by this tumor model, it is worth mentioning the ability to perform comparative biomechanical and transcriptomic analyses of confined and released cancer cells, enabling the identification of markers regulated immediately after intravasation. Since these studies are carried out in the absence of cytokines or chemoattractants, the experimental system more closely resembles the biophysical conditions present in primary tumors. Furthermore, this platform allows parallel investigation of these processes in non-malignant cells, such as fibroblasts and endothelial cells, as demonstrated in the present work.

Based on these facts, this study is focused on profiling the mechanotranscriptomic signatures of MCF-7 breast cancer cells when released from polymer-based artificial microtumors, mimicking the process of intravasation.

## 2. Results and Discussion

In human patients, cancer progression is characterized by multiple stages during which cells acquire a wide spectrum of features required to survive, proliferate, invade new tissues, and evade immunosurveillance. Among these stages, intravasation describes the release of cancer cells from a confined, semi-solid microenvironment into a soluble milieu, free of mechanical stress (i.e., solid stress).

To study this process in vitro, MCF-7 breast cancer cells were encapsulated within artificial microtumors, and their transcriptomic and biomechanical activities were compared with those of cells cultured on conventional flasks, as well as with cells freely released from these tumor-like matrices through a process mimicking intravasation (Figure 1A). As previously reported by our group, these artificial microtumors are characterized by displaying an elasticity of approximately 20 kPa (measured as Young’s modulus), consistent with the mechanical properties of breast tumors at metastatic stages.(^23,24^) Furthermore, the chemical composition of these matrices, made of alginate and gelatin (i.e., unraveled collagen), allows cancer cells to enzymatically degrade these scaffold, enabling their migration out of the matrix.

**Figure 1:**
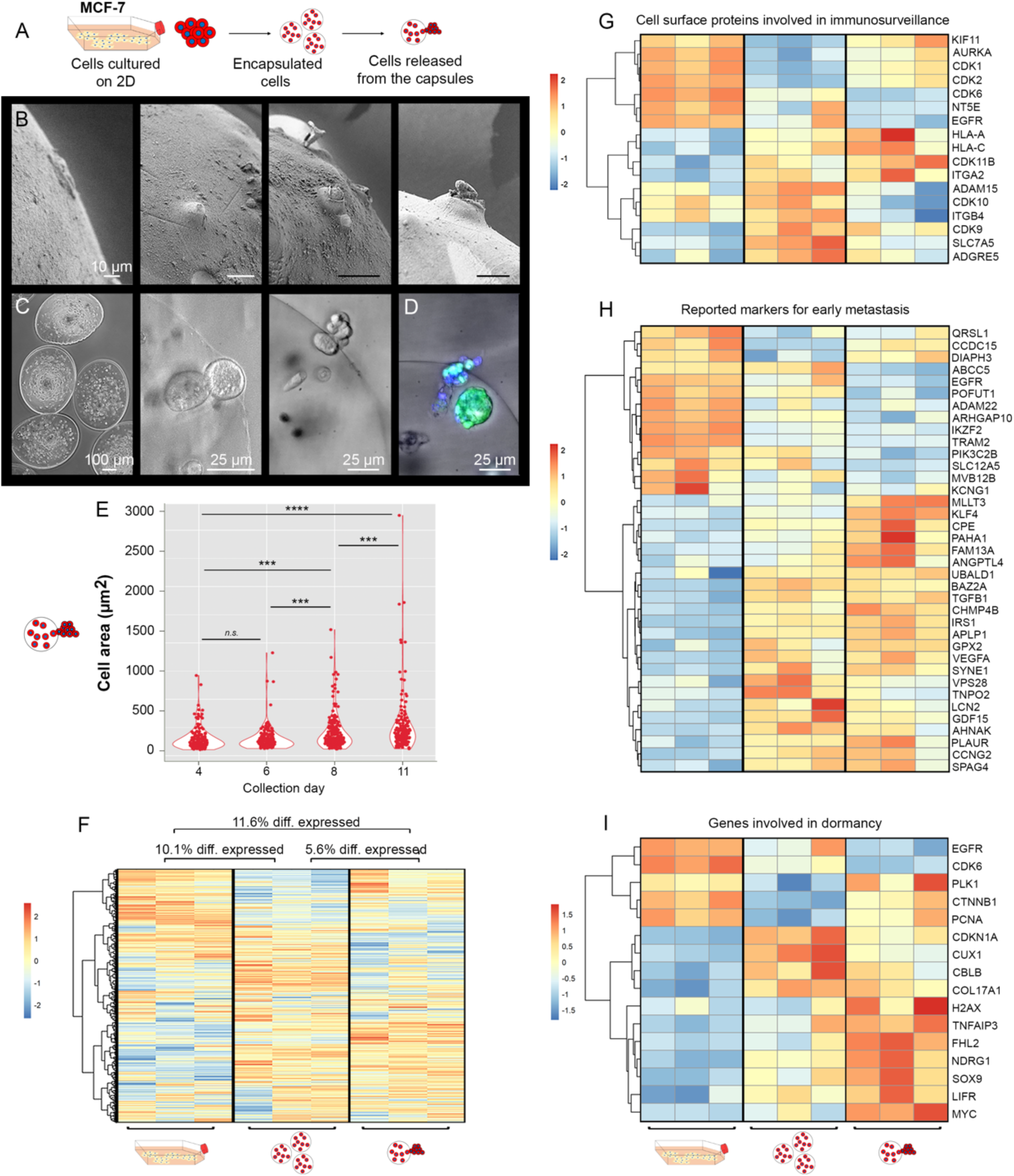
(A) Experimental design used in the preliminary phase of this work. In this study, cells were obtained from flasks (i.e., 2D), extracted from the bulk of the artificial tumors (i.e., after capsule dissolution), and characterized after their release from these scaffolds (i.e., intravasated-like cells). Images (B) and (C) show scanning electron microscopy and confocal microscopy images of MCF-7 cells in the process of being released from the artificial microtumors (i.e., intravasation-like process). Image (D) shows a cluster of cells during intravasation-like from the artificial microtumors. In that picture, the green color shows the cytosol of living cells (i.e., Calcein-AM), while the blue color shows the cell nuclei (stained with Hoechst). Plot (E) illustrates differences in the adhesion surface of MCF-7 cells that are released from artificial tumors (i.e., intravasation-like) at different time points after encapsulation. The heatmap in (F) displays differences in the full transcriptome of MCF-7 cells cultured under the three conditions described above, while (G), (H), and (I) show the differential transcriptomic expression of genes involved in immunosurveillance, early metastasis, and dormancy. In Figures (F), (G), (H), and (I), the side bars represent the z-score for each dataset.

Imaging of the intravasation-like process (Figures 1B, 1C and 1D) reveals that leading intravasating cells are subjected to intense shear stress and frequently die during this process. Noteworthily, the presence of naked and compacted nuclei can be observed on the surface of the scaffold, while viable cells remain within the artificial tumor (Figure 1D). These observations suggest that the cell release (i.e., intravasation-like) may involve a sacrificial mechanism, whereby leading cancer cells use their compact nuclei as a piston,(^8^) exerting forces against the boundary of the artificial tumors. As a consequence, a generation of pores and/or tracks is observed on the tumor surface, subsequently facilitating the escape of additional cells, while the leading cells themselves may die during the process.(^25^) This interpretation is supported by our previous studies, demonstrating that prior cells intravasate, a strong nuclear compaction is observed.(^20^)

In these artificial microtumors, cell release typically begins three days after encapsulation. Although the morphological characteristics of intravasated-like cells were not analyzed in detail in the present study, we evaluated the projected surface area of released cells after they seeded and attached to flat substrates adjacent to the microcapsules (i.e., 6-well plates) (Figure 1E). Our observations revealed the appearance of progressively larger cells after intravasation-like, suggesting the time-dependent emergence of distinct morpho-mechanical clones driven by biomechanical stress.(^26^) According to our previous publications, the increase in size and/or anchorage area can be due to changes in the cytoskeleton organization, as well as the generation of Polyploid Giant Cancer Cells (PGCC).(^22^) Based on these results, we decided to evaluate the mechanotranscriptomic activity of early intravasated-like cells, namely, those released at day 5 post cell immobilization.

Comparative transcriptomic analysis of cells extracted from the different culture conditions revealed that the largest differences were observed between cells cultured on 2D flasks and those released from the tumor-like microcapsules (11.6%), followed by differences between cells cultured on 2D and MCF-7 confined within microcapsules (10.1%). In contrast, cells confined within artificial microtumors and those collected from the polymer-based scaffolds exhibited the highest degree of similarity, with only 5.6% differential expression (Figure 1F).

To determine how mechanical stress affects the transcriptomic activity of cancer cells, we focused our subsequent analyses on genes involved in immunosurveillance, early metastasis, and cell dormancy. These genes were chosen, based on several publications in the field, as described in the supplementary tables 1, 2 and 3. Regarding surface proteins associated with immunosurveillance (Figure 1G), we observed differential regulation of several key mediators of antitumor immunity. As our experiments show, intravasated-like cells downregulate the expression of CDK10, NT5E, EGFR, and ADAM15, among others. Additionally, intravasated-like MCF-7 cells upregulate transcripts associated with Major Histocompatibility Complex class I (HLA-A and HLA-C). These results suggest that cells released from artificial tumors may be capable of eliciting a strong immune response, in contrast to what has been reported for highly metastatic cancer cells.(^27–31^) This behavior may be however explained, by the dynamic nature of cancer cell immunosurveillance, which is thought to evolve from early stages of intravasation to later stages, once cells have entered the blood or lymphatic vessels, as described in previous studies.(^32–35^) Additionally, it has been reported that the enhanced ability of cancer cells located in primary tumors to recruit macrophages could promote malignant progression by facilitating tumor cell invasion and intravasation, as well as by contributing to angiogenesis through extracellular matrix degradation, ultimately enhancing the entry of cancer cells into the bloodstream.(^36–38^) This information is supported by recent publications showing that downregulation of CDK10,(^39^) NT5E,(^40^) EGFR,(^41,42^) and ADAM15(^43^) may recruit macrophages, which could promote cancer progression.

After evaluating the expression of immunosurveillance-related proteins, we next focused our study on markers associated with early metastasis, including those regulated by hypoxia, nuclear repair, and early cell dissemination (Figure 1H). The full list of analyzed genes, as well as original publications describing them can be found in the tables 1, 2 and 3. Analysis of these malignant hallmarks revealed that cells cultured on 2D surfaces differed markedly from those confined within or released (i.e., intravasated-like) from 3D microtumors. Among the markers upregulated in cells cultured in, or released from 3D environments, we detected genes involved in intravasation-like (i.e., KLF4, PLAUR),(^44–45^) early metastatic dissemination (i.e., LCN2, AHNAK, BAZ2A, APLP1, ANGPTL4),(^46–50^) nuclear rupture and repair (i.e., SYNE1, SPAG4),(^51,52^) and hypoxia-associated pathways (i.e., P4HA1, FAM13A, VEGFA).(^53–55^) Notably, SPAG4,(^56^) P4HA1,(^57,58^) and BAZ2A(^59^) have been additionally described as pan-cancer hallmarks, having a relevance which surpasses breast cancer pathologies, targeting other types of malignant pathologies characterized by the existence of solid tumors.

Further, we evaluated the role of mechanical stress in regulating cancer cell dormancy. The full list of analyzed genes, as well as original publications describing them, can be found in the supplementary tables 1, 2 and 3. As shown in Figure 1I, cells cultured within or released from artificial microtumors exhibited a clear upregulation of dormancy-associated markers, compared to cells cultured on 2D flasks. In particular, increased expression of SOX9(^60,61^) and H2AX(^62^) was observed, with special attention given to MYC(^63^) and FHL2.(^64^) In particular, MYC upregulation has been shown to tune the STING pathway, thereby controlling cancer cell dormancy during DNA repair, while simultaneously modulating inflammatory signaling in breast cancer cells.(^63^) In parallel, FHL2 has been reported to be upregulated in cancer cells cultured in stiff 3D environments,(^64^) where it regulates dormancy in response to mechanical load. This finding reinforces the idea that cancer cells preconditioned to stiff 3D milieus exhibit a broader array of hallmarks associated with progression and malignancy compared with cells cultured in very soft 3D environments, or even on 2D surfaces.

In the next phase of this research, we compared the regulation of transcripts involved in cell biomechanics (i.e., YAP/TAZ) and the expression of catalytic enzymes, such as metalloproteinases (MMPs) and A Disintegrin and Metalloproteases (ADAMs) (for details, please check the supplementary tables 4 and 5 respectively). As Figure 2A shows, the YAP/TAZ pathway is strongly upregulated in released cells and, to a lesser extent, in cells cultured within the artificial tumor, followed by cells cultured on 2D flasks. These results suggest that cancer cells exhibit enhanced biomechanical activity after exposure to solid stress. Similar analyses were performed for catalytic enzymes involved in extracellular matrix degradation (i.e., MMPs and ADAMs), revealing that these proteins are most strongly upregulated in confined cells, followed by intravasated-like cells from artificial microtumors, and finally cells cultured on flat surfaces (Figure 2B). This trend is consistent with the expectation that invasive cells increase the synthesis and secretion of catalytic proteins. Regarding specific ADAM family members, ADAM15 is notably upregulated in cells within artificial microtumors, a finding consistent with its established role in basement membrane disruption, prior to cells undergoing intravasation in vivo.(^65–68^) Additionally, it is worth noting that the strongest upregulation of MMP1 and MMP13 is detected in released cells. With regards to MMP1, this enzyme has a strong collagenase activity and it drives tumor invasion, metastasis, and angiogenesis by degrading the extracellular matrix, which is consistent with the chemical composition of the tumor and the invasive activity of MCF-7 cells.(^68^) On the other hand, the presence of MMP13 could play a role as part of the features displayed by cancer cells during the expression of mechanical memory, which is referred to the probability shown by cancer cells cultured in stiff tumors, to metastasize into hard tissues, such as bone.(^69,70^) Since MMP13 mediates osteoclast differentiation and bone colonization, the upregulation of this enzyme may be a preliminary mechanisms involved in osteocolonization.(^71^)

**Figure 2:**
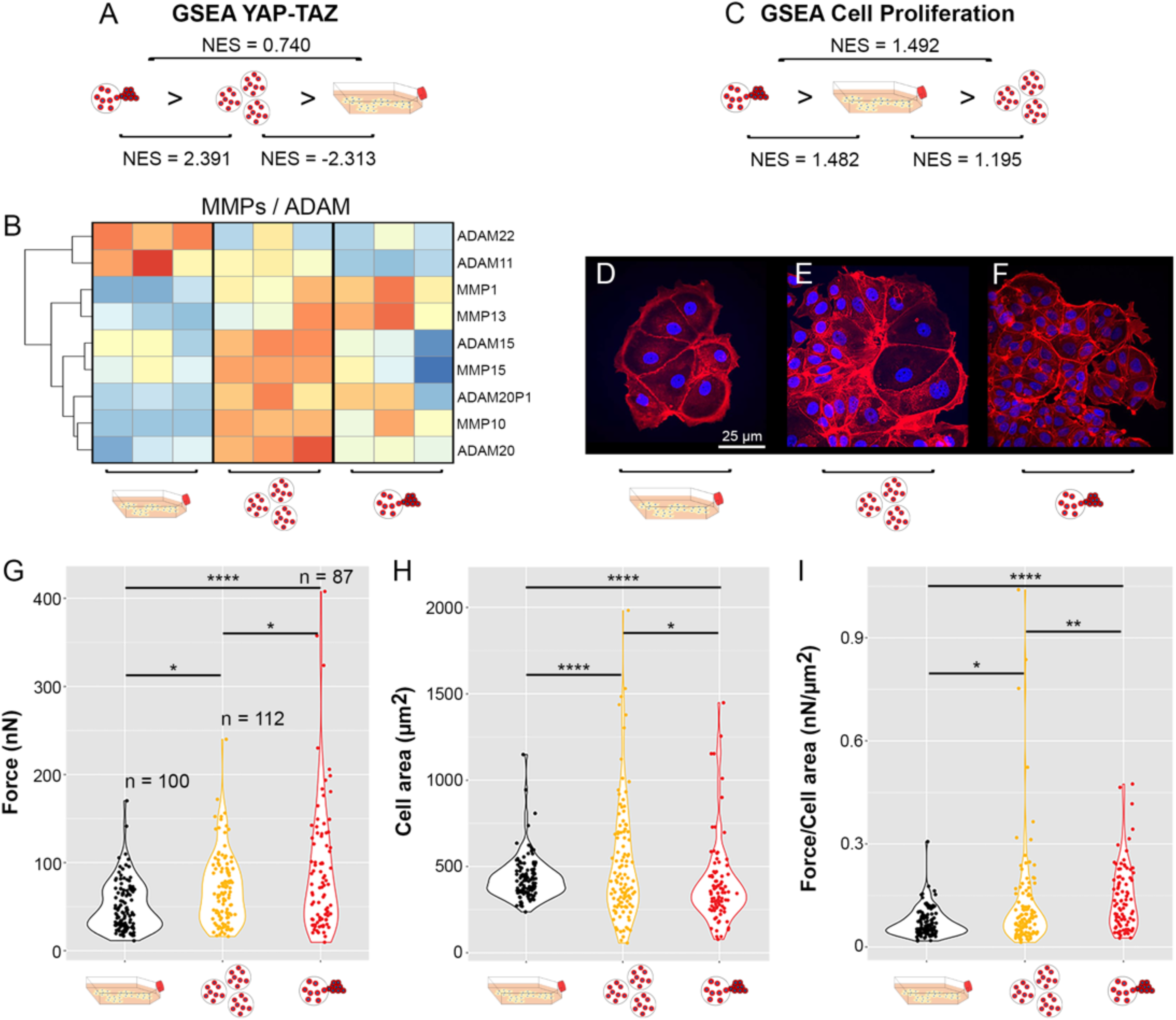
Transcriptomic and biomechanical analysis of cells cultured on flasks, extracted from the artificial microtumors and released from these scaffolds. Images (A) shows the comparative GSEA analysis of transcripts involved in the YAP/TAZ pathway. The heatmap (B) displays the transcripts related to the expression of MMPs and AMDAs in MCF-7 cells cultured in each milieu. GSEA results in (C) indicates that cells released from the capsules shows a higher proliferative capability compared to cells cultured on flat surfaces and those entrapped in the artificial microtumors respectively. Images (D), (E) and (F) depict the morphology of MCF-7 cells cultured on flasks (D), extracted from the artificial microtumors (E) and released from these scaffolds (F). In these images, the cell cytoskeleton is marked in red (i.e., actin fibers; stained with rhodamine-phalloidin), and the nuclei in blue (stained with Hoechst). Images (D), (F) and (G) have the same scale. Plots (G), (H) and (I) show the results of single-cell traction force microscopy analysis carried out in cells cultured/extracted in their respective milieus during/after 5 days of culture. Graphic (G) shows the total forces exerted by cancer cells, while the plot (H) displays the anchorage area evaluated in studied cells. Finally, the graphic (I) shows the ratio Force/Area for single cell. In these plots, comparative studies were carried out by using the Kolmogorov–Smirnov test. In Figure B, the side bars show the “z-score” for each study.

**Figure 3:**
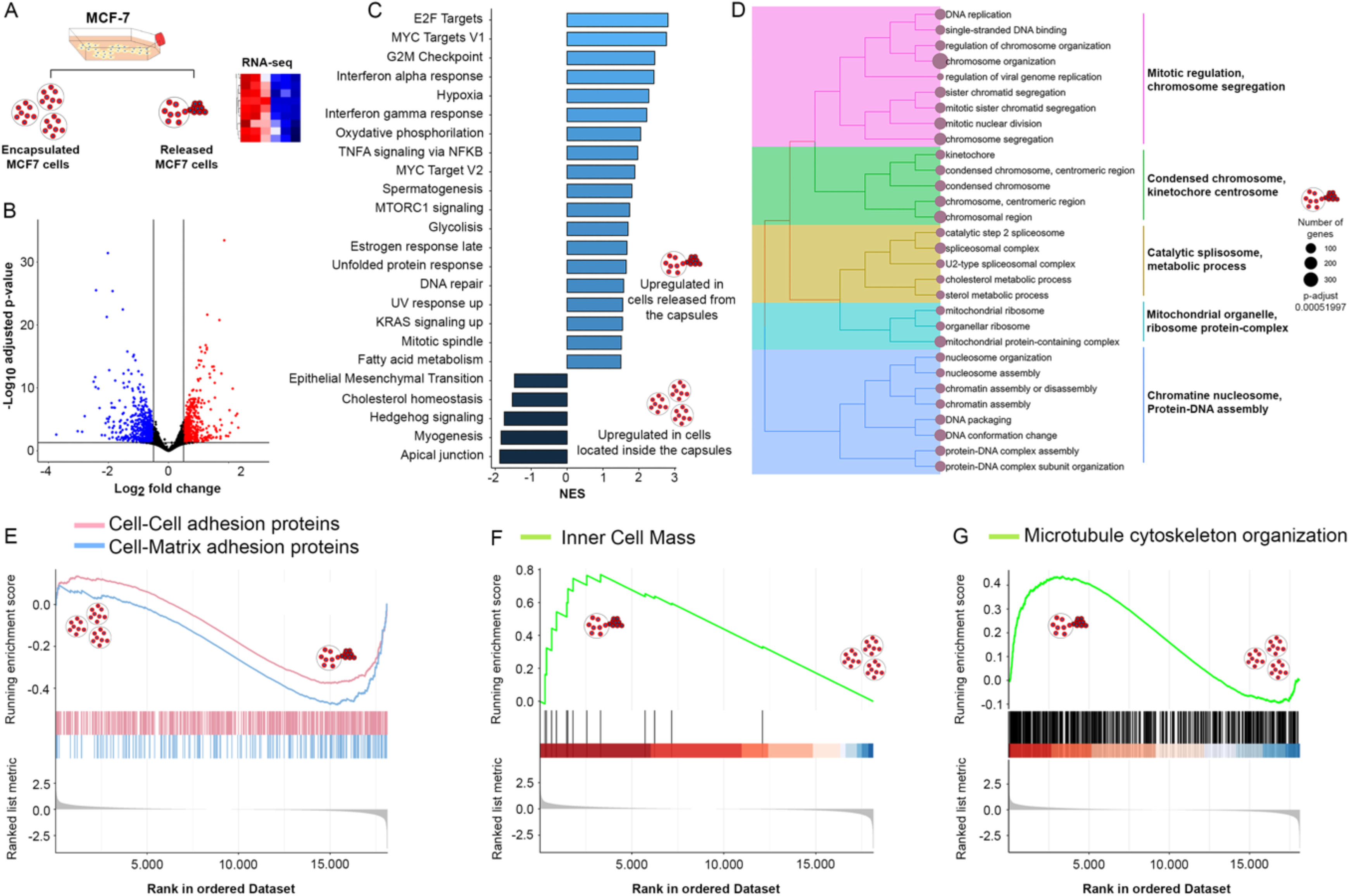
(A) Experimental design used to compare the transcriptional activity of cells cultured inside artificial microtumors and those released from these tumor-like scaffolds. (B) Volcan plot showing the differential transcriptional activity of both analyzed cell types. In this experiment, the number of differentially upregulated genes is 588, while the number of downregulated genes corresponds to 670. (C) Differential tuning of malignant pathways in confined and released cells in/from the artificial microtumors. (D) Specific biological and/or metabolic markers expressed in intravasated-like cells. GSEA analysis of transcripts involved in cell-cell and cell-matrix adhesion (E), Inner cell mass (F) and Microtubule organization (G).

**Figure 4:**
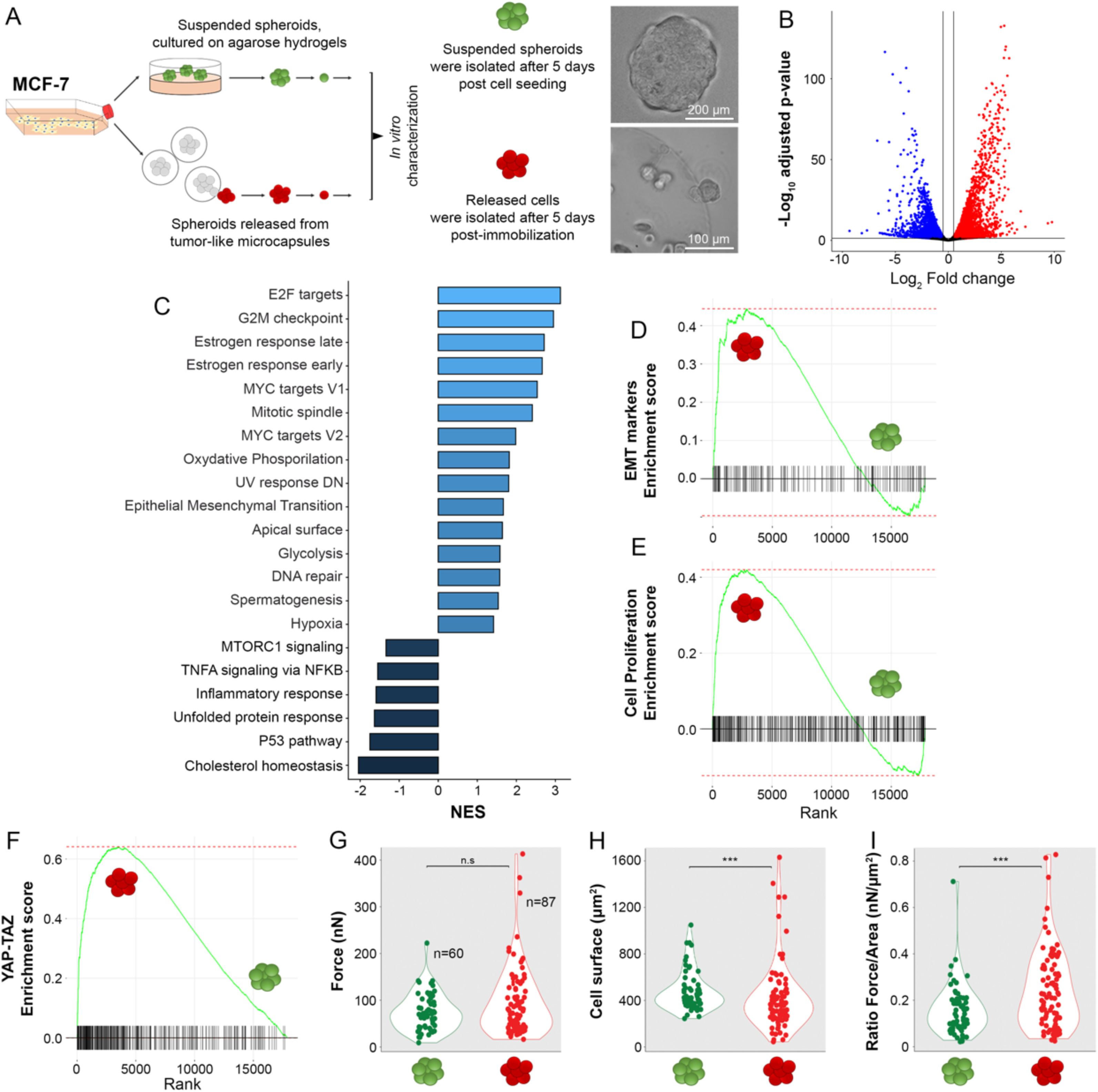
(A) Experimental design used to compare MCF-7 cells cultured as suspended spheroids (green) with cells released from artificial microtumors (red). The images on the right show representative spheroids from each condition. (B) Volcano plot showing the differential transcriptional activity between the two analyzed cell types. In these experiments, 4091 genes were significantly upregulated, while 3441 genes were downregulated. (C) Differential regulation of malignant pathways in confined and released cells within/from artificial microtumors. (D), (E) and (F) show Gene Set Enrichment Analysis (GSEA) of epithelial-mesenchymal transition (D) proliferation (E) and YAP/TAZ pathway. (G), (H), and (I) illustrate the biomechanical activity of MCF-7 cells analyzed by single-cell traction force microscopy. Statistical comparisons were performed using the Kolmogorov–Smirnov test.

The subsequent Gene Set Enrichment Analysis (GSEA) was focused on assessing the transcript expression levels of proliferation-related genes in MCF-7 cells (Figure 2C). Our results indicate that cells released from artificial microtumors exhibit the highest expression of proliferation-associated markers, followed by cells cultured in flasks, whereas cells maintained within artificial microtumors show the lowest expression levels. These results suggest that cells actively respond to the presence of solid stress by enhancing their proliferative capabilities, results which are consistent with our previous publications in the field.(^72^) We further hypothesize that these genes are predominantly expressed once intravasated cells encounter a tissue that allows them to anchor and spread (as shown below, in Figure 5C). In contrast, while circulating in blood or lymphatic fluids, these cells likely remain in a quiescent state.

**Figure 5:**
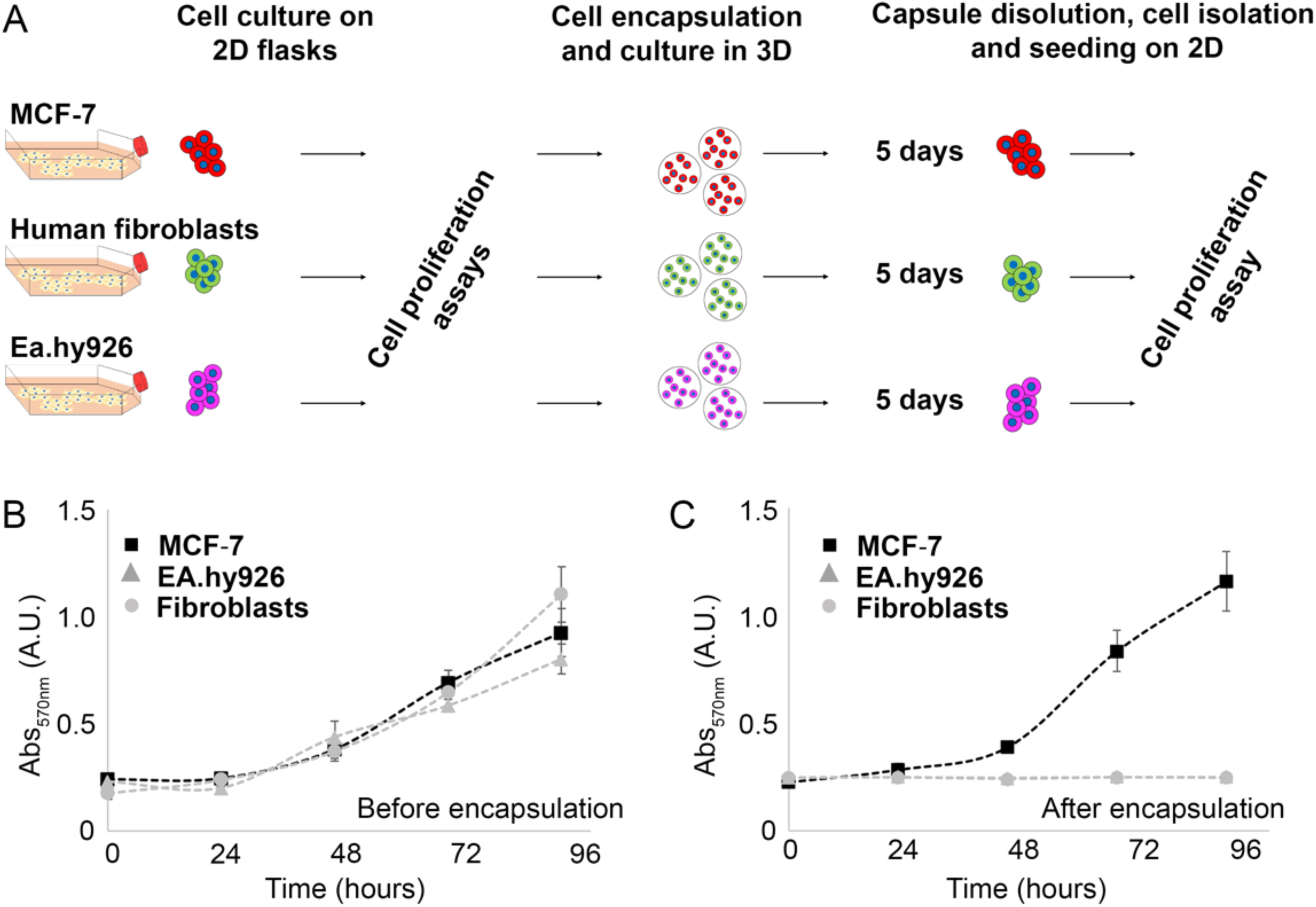
(A) Experimental design used to evaluate the role played by the biomechanical stress in the proliferative capabilities of MCF-7 breast cancer cells, Human fibroblasts and endothelial cells (Ea.hy926). Plots (B) shows the proliferation profile of MCF-7 breast cancer cells (▪), Human fibroblasts (⬤) and endothelial cells (▴) prior encapsulation, while (C) displays duplication profile of same cells after being cultured in the artificial tumors during 5 days.

To determine how confinement and mechanical stress influence the morpho-mechanical behavior of cancer cells, we characterized MCF-7 cells by confocal microscopy (i.e., morphological cues) and by single-cell traction force microscopy (TFM). In this work, TFM was using to assess the biomechanical activity of cancer cells pre-cultured in different milieus. Briefly, in this method, cells are seeded onto soft polyacrylamide hydrogels embedded with fluorescent microbeads. As cells exert traction forces, the beads are displaced, allowing their movement to be tracked and enabling quantification of the biomechanical activity of adherent cells, as well as their morphological characteristics (i.e. cell-matrix anchorage area). As shown in Figures 2D to 2F, cells cultured on flat surfaces display relatively homogeneous morphology and size compared with cells cultured in 3D. Furthermore, our results show that cells released from artificial microtumors exhibit higher biomechanical activity than cells cultured in the other two experimental conditions (Figure 2G), while also displaying a lower average cell area (Figure 2H). Overall, cells released from confinement are considerably more heterogeneous than those cultured within artificial tumors or on flat surfaces (Figure 2I). These findings suggest that small and capable to exert higher magnitudes of mechanical forces are preferentially released from the microtumors, potentially increasing their metastatic potential.

After performing comparative analyses of aforementioned cells, we focused our study on evaluating transcriptomic differences between those confined within microtumors and MCF-7 released from confinement (Figures 3A and 3B). The main objective of these assays was to determine the similarities and differences between these two populations, and to identify differentially expressed malignant markers associated with pathological progression. As shown by GSEA analysis (Figure 3C), extravasated MCF-7 cells exhibit a stronger pathogenic response compared with cells remaining inside the microtumors. This is exemplified by the upregulation of markers associated with malignant pathways, including E2F targets,(^73–75^) MYC targets V1,(^73–75^) and MYC targets V2,(^73–75^) as well as pathways related to hypoxia,(^76^) and DNA repair.(^77^) Metabolically speaking, released cells also upregulate markers of glycolysis, fatty acid metabolism, and oxidative phosphorylation (Figure 3C).(^78^)

With respect to the upregulation of EMT markers in confined cells compared to intravasated-like cells, we hypothesize that this process is necessary to enhance the migratory capacity of cancer cells when located within a confined and semi-stiff microenvironment. However, once released, these cells appear to enter a dormant state (Figure 1I and Figure 3F), allowing them to repair damaged cellular structures (i.e., DNA)(Figure 3C and 3D), to evade immunological surveillance (Figure 1G), and to facilitate the colonization of distant tissues, away from the primary tumor site, as discussed below.

These observations are supported by the upregulation of G2M checkpoint and MTORC1 hallmarks, as well as pathways involved in oxidative phosphorylation, which have been reported to play a key role in DNA repair and in the transition of stem cells from quiescence to a proliferative state.(^79,80^) Additionally, these findings align with the expression of specific markers involved in mitotic regulation and chromosome organization (Figure 3D), strongly suggesting that mechanical stress stimulates cancer cells to enter dormancy, likely because they require time to repair damaged DNA, which could be caused by shear stress, most probably occurring during cell release, as already reported by other teams.(^51,52^)

Interestingly, after intravasation-like, released cells show a decreased expression of proteins involved in cell-matrix and cell-cell adhesion (Figure 3E). Intravasated-like MCF-7 also maintain an undifferentiated state, which is exemplified by the capability of cancer cells to keep their inner mass under hypoxic and glycolytic conditions (Figure 3F), a phenomenon traditionally reported in pluripotent stem cells, cancer stem cells, and pre-implantation embryos prior to differentiation.(^81–83^) These findings suggest that small clusters of cells within the microtumors disaggregate during release and undergo dedifferentiation, facilitating migration and subsequent invasion of tissues distant from the primary tumor, mediated by the increment in their biomechanical activity, as previously shown in Figure 2H. Furthermore, as shown in Figure 3G, when cells are cultured in 3D environments, these processes appear to be mediated by the organization and/or repair of microtubules, which could act as a mechanosensors, as already reported by several groups.(^84–86^)

When MCF-7 cells migrate out of the artificial microtumors, they are released as small clusters, typically composed of five to ten cells (data not shown). Based on these observations, in the next series of analysis we compared the expression of malignant markers in cells released from artificial microtumors with those expressed in cell aggregates cultured on non-adherent surfaces (Figure 4A). These spheroids mainly differ in the magnitude of mechanical stress experienced by the cancer cells, which is mostly driven by neighboring cells. Yet, since suspended aggregates primarily sense the elasticity of neighboring cells,(^87^) they are subjected to a lower mechanical load compared with cells cultured within tumor-like scaffolds.

Comparative transcriptomic assays of cells cultured as spheroids versus intravasated-like cells revealed clear differences (Figures 4B and 4C). Notably, released cells upregulate markers associated with pathogenicity, including E2F, and MYC targets V1 and V2. From a metabolic perspective, hallmarks of hypoxia and glycolysis were also upregulated (Figure 4C), indicating that intravasated-like cancer cells express a broader set of proteins and enzymes that may enhance survival under hostile environmental conditions and support persistence during dormancy.

Previous studies were complemented by GSEA assays of three main pathways involved in cancer pathogenicity, namely EMT, cell proliferation, and biomechanical activity (i.e., YAP/TAZ) (Figures 4D to 4F). In all three cases, released cells exhibited increased expression of these transcripts compared with spheroid-forming cells (i.e., cultured on non-cell adhesive hydrogels), displaying an enhanced malignant potential. With the purpose to experimentally validate the upregulation of YAP/TAZ hallmarks, we carried out single-cell traction force microscopy assays on these cells. As shown in Figures 4G to 4I, released cells exert stronger cell-matrix attachment forces while displaying a smaller adherent area, resulting in subpopulations with a higher Force/Area ratio (i.e., smaller cells with increased attachment strength). These cells also exhibited greater morpho-mechanical heterogeneity. Together, these results demonstrate that cells released from artificial microtumors, compared to cells cultured as spheroids, more closely recapitulate the biological patterns observed in cells intravasated from primary tumors, compared to cells cultured as conventional spheroids, including, but not restricted to their capability to repair damaged structures (Figure 4C) and to attach and colonize foreign tissues (Figures 4G to 4I), corroborating preliminary experiments performed by our teams, comparing the activity of cells cultured as 3D spheroids with those located inside the artificial microtumors.(^20^)

In the final stage of our study, we evaluated the invasive behavior of cancer cells relative to that of healthy cells. The experimental design consisted of two sequential steps. First, RFP-derived MCF-7 breast cancer cells, GFP derived human fibroblasts, and human endothelial cells (EA.hy926 cell line, stained with cell mask) were cultured separately within artificial tumors (monoculture) (Figure 5A). The aim of these experiments was to assess their proliferative capacity before and after exposure to mechanical stress. As shown in Figure 5B, prior encapsulation, all three cell types proliferated on 2D surfaces with similar duplication rates. However, after five days of cell confinement, only RFP-derived MCF-7 cells were able to recover their proliferative capabilities (Figure 5C), while GFP-human fibroblasts and endothelial cells seems to enter in a senescent stage.

To further test whether confinement and mechanical stress affect invasive behavior, the three cell types were then co-cultured within the microtumors, at a ratio of 1:3:3 (MCF-7 / fibroblasts / endothelial cells) (Figure 6A). After five days of co-culture, only MCF-7 cells were able to proliferate, migrate, and escape from the capsules, whereas healthy cells remained trapped within the artificial tumors and few cells were detected outside the capsules (Figures 6B and 6C). These results support the idea that mechanical stress promotes malignant responses that are not exhibited by healthy cells. Furthermore, these experiments additionally demonstrate that the artificial tumor platform described here could serve as an effective strategy to enrich and isolate highly malignant cancer cells from complex multicellular samples, based on the differential sensitivity of healthy and neoplastic cells to mechanical stress.

**Figure 6:**
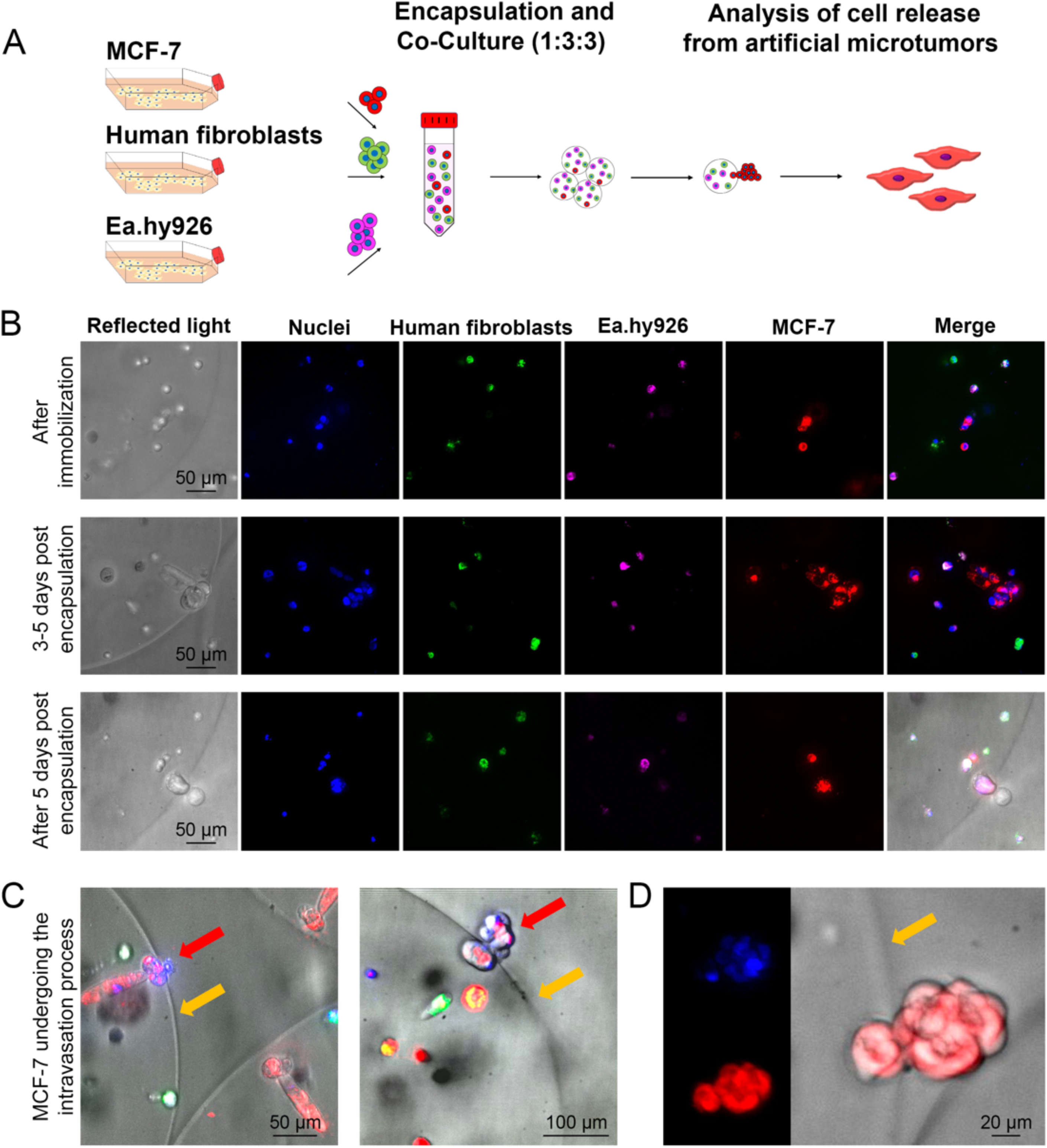
(A) Experimental design used to evaluate the migration and release of RFP-derived MCF-7 breast cancer cells, GFP-derived human fibroblasts and endothelial cells (stained with cell mask), when co-cultured in artificial tumors for 5 days. (B) show the results of these experiments. In these pictures, the blue color shows the staining of cell nuclei, the green fluorescence corresponds to the presence of human fibroblasts, human endothelial cells were stained in magenta, while RFP-derived MCF-7 cells are shown in red. Images (C) and (D) show the release of RFP-derived MCF-7 cells from tumor-like microcapsules. In these images, cells undergoing intravasation-like are indicated by red arrows, while the boundary of the artificial microtumor is marked by orange arrows. Figures (C and D) show a of RFP-derived MCF-7cells in process of intravasation-like.

## 3. Conclusions

Intravasation represents the earliest stage of cancer metastasis. This process is characterized by the release of malignant cells from a semi-solid three-dimensional environment (i.e., primary tumors), into an aqueous milieu through an impaired vessel wall (i.e., blood or lymph vessels). Primary breast tumors microenvironments are made of crosslinked extracellular matrix, typically dense and poorly degradable. These biophysical characteristics restrict migration and proliferation of entrapped cells and subjecting them to limited oxygen and nutrient supply. Because studying intravasation-like in vitro is extremely challenging, in this work we cultured MCF-7 breast cancer cells within a new tumor model known as artificial microtumors, with the purpose to study their release (i.e., intravasation-like process), mediated by metalloproteinase expression and biomechanical forces.(^20^)

Based on these facts, this study was focused on characterizing the mechanotranscriptomic profile of cancer cells under different culture conditions, including cells seeded on 2D flasks and those cultured as 3D spheroids. The outcomes of this research, namely the mechano transcriptome differences between cells cultured on 2D, those confined in artificial microtumors, and MCF-7 cells released from these scaffolds, are summarized in Figure 7. As illustrated, the shift in mechanical stress experienced by released cells, combined with increased availability of nutrients and oxygen, triggers upregulation of multiple malignant hallmarks, including proliferation, YAP/TAZ activation, enhanced biomechanical activity, DNA repair, and the expression of markers for cell dormancy. At the same time, these cells show reduced capacity for adhesion to surrounding tissues and decreased expression and secretion of catalytic enzymes such as MMPs and ADAMs. We hypothesize that these features may arise for two main reasons: (i) to optimize energy use, and (ii) to increase the likelihood that cancer cells remain suspended in the bloodstream or lymphatic circulation for an undefined period of time, thereby enhancing their capacity to disseminate and colonize distant organs rather than tissues adjacent to the primary tumor.

**Figure 7:**
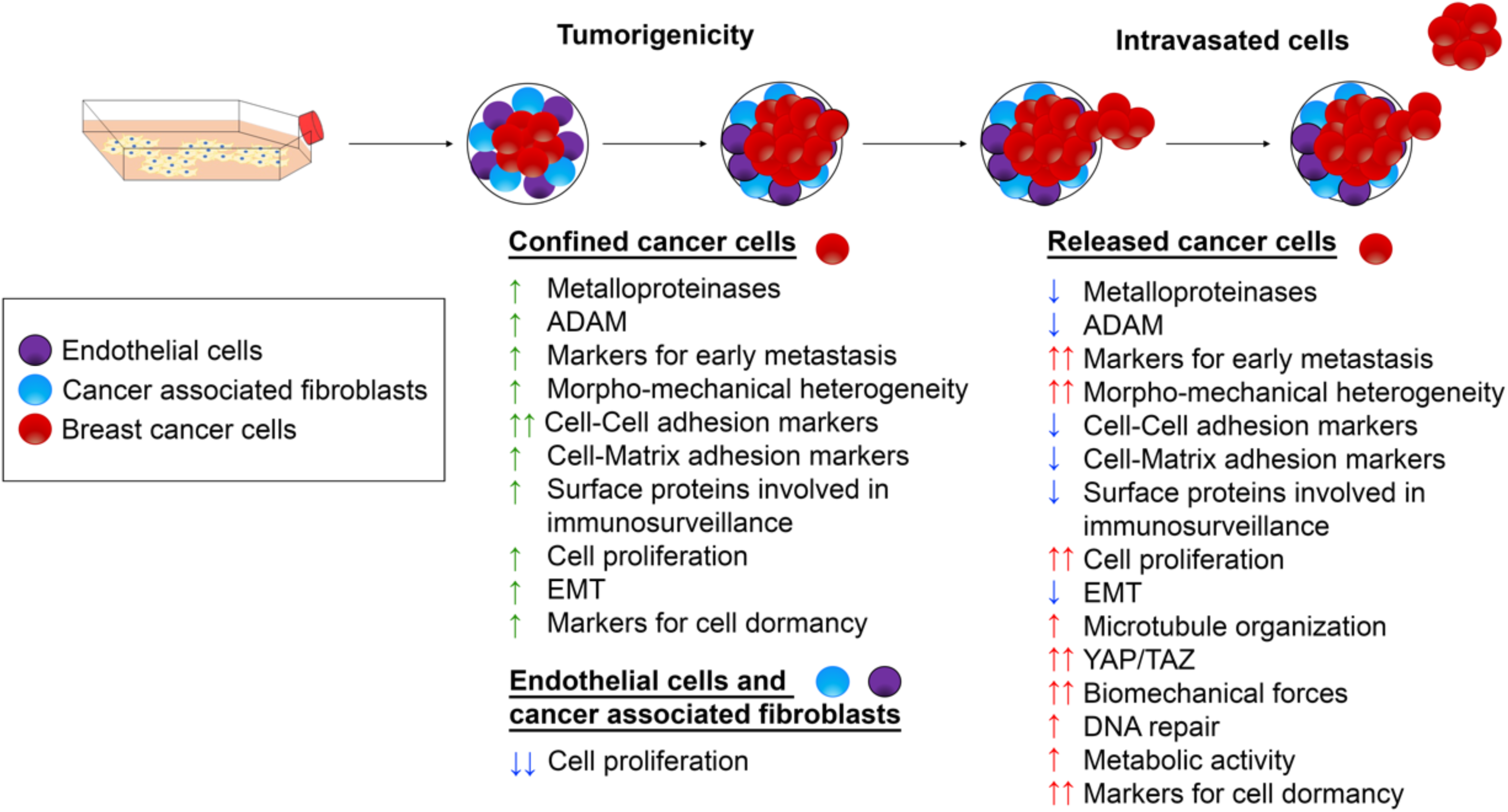
Summary of the results obtained in this study. Green arrows indicate malignant hallmarks that are upregulated in cells located within artificial tumors compared to cells cultured on 2D flasks. Red arrows represent pathways upregulated in intravasated-like breast cancer cells relative to their counterparts confined within tumor-like microcapsules. Finally, blue arrows denote downregulated neoplastic pathways, either in endothelial cells and/or cancer-associated fibroblasts compared to MCF-7 cells cultured within the same capsules, or in intravasated-like MCF-7 cells with respect to cells remaining inside the artificial microtumors.

Additionally, it is worth noting the upregulation of hallmarks described in dormant cells. We believe that this quiescent period allows breast cancer cells to repair DNA damage and prepare for subsequent invasion of new tissues. Interestingly, this behavior is not observed in healthy cells, which enter a senescent state when subjected to mechanical confinement. Regarding immunosurveillance markers, our results additionally suggest that early intravasated-like cells may be more detectable by the immune system, likely due to upregulation of major histocompatibility complex class I genes (HLA-A and HLA-C). This behavior can be explained by the possibility to attract macrophages, which could degrade the tumor matrix, thus enabling the release of neoplastic cells. Finally, because released cells from the artificial microtumors also downregulate cell-cell adhesion markers, the presence of single cells or very small aggregates may reduce the probability of immune detection, as they circulate in patient fluids, increasing the probabilities of surviving and metastasis, processes which however, require the mechanical stress to be triggered.

Finally, with the information described in this paper, we created a chart, comparing the differences in malignant marker for cancer located inside the artificial microtumors and those released from these tumor-like scaffolds. As Figure 7 shows, the 5.6 % of transcriptomic difference already described between these two cell types (Figure 1F) is extremely relevant for the pathogenic activity of cancer cells, which display and tune a series of hallmarks and pathways required to foster the malignant progression, information which could be extremely relevant to generate differential therapies, which could be either targeting cells located inside tumors, or those already released from the neoplastic niche.

## 4. Materials and Methods

### 4.1 Cell culture

The breast cancer cell line MCF-7 (DSMZ Cat. N° ACC 115) was sourced from our departmental cell bank. Additionally, we used MCF-7 stably expressing RFP (Cat. No. T3932, Abm) in experiments described in figure 5 and 6. Human mammary fibroblasts expressing GFP,(^88^) were generously provided by Dr. Charlotte Kuperwasrer (Tufts University, MA). EA.hy926 endothelial cells were purchased at ATCC (CRL-2922™). For experiments shown in Figures 5 and 6, cells were stained using cell far-red Mask (HCS CellMask™, Invitrogen, Germany). All cells were cultured in DMEM cell culture media (High glucose, Gibco, USA), supplemented with 10% v/v of FBS (Fetal Bovine Serum; Corning, USA) and 1.0% v/v penicillin-streptomycin (Gibco, USA) and 0.1% v/v gentamycin (Sigma-Aldrich).

### 4.2 Preparation of suspended spheroids and tumor-like microcapsules

Suspended spheroids were obtained by seeding 20·10^3^ cells in 24 well plates previously coated with 250 µl of agarose (1.0% w/v) (Merck, Germany), following published methods.(^20^) Tumor-like microcapsules were made using a micro-encapsulator device under patenting process (EP23207537.4),(^89^) as well as a microencapsulator Büchi E-390.(^20^) Microcapsules were made of 1.0% w/v alginic acid (Sigma-Aldrich, Germany) and 1.0% w/v gelatin type A (Sigma-Aldrich, Germany). These polymers were dissolved in a solution containing 0.1M HEPES (pH 7.4, Sigma-Aldrich, Germany), 1.0 % w/v NaCl, and 1.0 % v/v penicillin-streptomycin (Sigma-Aldrich, Germany). This blend was then filtered by using a 0.45 µm pore diameter filter (Merck Millipore, Germany), and loaded with MCF-7 cells with a density of 5·10^5^ cells/mL. Alternatively, co-culture cell encapsulation was performed by mixing the hydrogel with the cell mixture in a 1:3:3 ratio (Cancer/Fibroblasts/Endothelial cells), at a density of 1x10⁶ cells/mL.

To crosslink the alginate-gelatin microcapsules, we used a solution of 0.6 M of CaCl2, dissolved in 1.0% w/v HEPES (Sigma-Aldrich, Germany) containing 1.0% w/v NaCl (pH 7.4).

### 4.3 Isolation of MCF-7 released from artificial microtumors

To isolate intravasated-like cells, artificial microtumors (loaded with MCF-7 breast cancer) cells were placed on cell strainers (Corning, Germany) with a 100 µm pore size. The strainers were then located in 6-well plates coated with a thin layer of agarose (1.0% w/v; Merck, Germany). To isolate cells released on day 5 (post-encapsulation), microtumors were placed on the strainers on day 4 (post-cell immobilization) and removed on day 6 (post-cell immobilization), allowing the collection of floating cells (i.e., which have been released on day 5) for subsequent transcriptomic analyses.

For the evaluation of the adhesion surface of released cells, the same strategy was applied to characterize cells released on days 4, 6, 8, and 11. In this case, however, the strainers were placed in standard adherent 6-well plates (not coated with agarose), enabling cells to attach and spread onto these surfaces.

### 4.4 Isolation of cells from alginate-gelatin microcapsules, and from 3D spheroids

MCF-7 cells were isolated from the microcapsules, by immersing them for 1 minute in a solution containing 22.5 mM of sodium citrate dihydrate, 60 mM EDTA (Sigma-Aldrich; Germany), and 150 mM of sodium chloride (Sigma-Aldrich; Germany, pH 7.4), following protocols already published.(^20^) After that, cells were isolated by centrifugation (1300 rpm, 4 min). Concerning the isolation of MCF-7 cells from the spheroids, these cell aggregates were initially resuspended in trypsin-EDTA (Pan Biotech, Germany), and placed in the incubator for 3 minutes. This procedure was followed by a 10 minutes incubation in a solution 1:1 of cell culture medium and trypsin-EDTA, followed by a centrifugation step (1300 rpm, 4 min) to recover single cells.

### 4.5 RNA-Sequencing assays and transcriptome analysis

RNA extraction was carried out according to protocols already published.(^20^) All experiments were carried out in triplicate. 75 bps paired- end sequencing was performed on a HighSeq 4000 Sequencer (Illumina) with an average number of 20 million reads per sample. After a quality check using FastQC v0.11.9, reads were aligned to the human reference genome (hg38) using the STAR alignment software (v 2.5.2a). After mapping, only reads that mapped to a single unique location were considered for further analysis. The mapped reads were then used to generate a count table using the feature-counts software (v 1.4.6-p5). The raw reads were filtered, normalized, and visualized by using R version 4.0.1. The differential expression analysis was done using the DESeq2 standard approach. Adjusted p-values were calculated using the Benjamini-Hochberg method within DESeq2. Differentially expressed genes with a log2FC of ±0.5 and an adjusted p-value <0.05 were considered statistically significant and visualized with ggplot2. The heatmaps were plotted with pheatmap (v.1.0.12) using a normalized variance-stabilizing transform (VST) in the DESeq2 framework.

### 4.4 Single Cell Traction Force Microscopy Assays

Single Cell Traction Force Microscopy (TFM) assays were performed following published methods.(^22^) Briefly, single cells were seeded on flat polyacrylamide gels coated with collagen type-I, following protocols previously described in literature. These gels had an elasticity of 8.5 ± 2.8 kPa (measured as Young’s Moduli). The elasticity of the polyacrylamide gels was analyzed by using a PIUMA nanoindentation device, as described in our previous publications. After cell attachment and spreading (overnight), forces exerted by single cells were determined using a fiji/MatLab-based software gently sourced by Dr. Martial Balland (Laboratoire Interdisciplinaire de Physique, Grenoble, France), and described in reference 22.

### 4.5 Cell imaging and staining

Cells were imaged using a Spinning Disc Axio Observer Z1 from Zeiss (Optical Imaging Competence Centre OICE). Immunostaining protocols were performed as follows: MCF-7 cells were isolated from the different milieus and seeded onto glass coverslips (22 X 22 mm). As next, cells were fixed using a 4.0% w/v paraformaldehyde, dissolved in PBS (Sigma-Aldrich, USA). Cells were then permeabilized for 15 min with 0.1% Triton X-100 and blocked for 30 min at RT with 5.0% fetal bovine serum FBS/PBS solution. This step was followed by their staining with Phalloidin-Rhodamine (Invitrogen, Germany), and Hoechst 33342, 1.0 μg/mL in PBS (Invitrogen, Germany), following the provider’s instructions.

### 4.6 Cell proliferation assays

Studies of cell duplication were carried out with MCF-7 cells isolated from 2D and 3D milieus, following published methods.^[20]^ Cell proliferation studies were performed using (3-(4,5-dimethylthiazol-2-yl)-2,5-diphenyl-2H-tetrazolium bromide (i.e., MTT; M6494, Thermofisher, Germany).(^90^) Initially, 5x10³ cells were seeded in a 96-well plate and incubated for different times (24, 48, 96, 120 hours). After each post-seeding time, the cells were incubated for one hour with MTT (1.0 mg/mL). Then, the formazan crystals were dissolved by adding 100 µL of DMSO:Isopropanol (3:2) (both purchased at Sigma-Aldrich; Germany). The absorbance was measured at 570 nm in a multi-plate reader (Tecan Infinite F200, New York, NY, USA), following protocols already reported by other research teams.(^90^) The results of three independent experiments were expressed as the average percentage relative to the initial time.

### 4.7 Gene set enrichment analysis and heatmap assays

The Gene Set Enrichment Analysis (GSEA) method was employed for the RNA-seq data involving 100.000 permutations of the following conditions (i.e., Encapsulated cells v/s Cells cultured on 2D flasks; Cells cultured as spheroids v/s Cells cultured on 2D flasks, and Encapsulated cells v/s Cells cultured as spheroids). A weighted enrichment score was utilized. Pre-ranked genes with an Entrez gene ID annotation were based on the DESeq2 and the fold difference between the two analyzed conditions. The Hallmark gene sets from MsigDB (v7.5.1) were used to perform GSEA via fgsea (v1.20.0). The respective gene dataset for immunosurveillance, early metastasis and dormancy are shown in the supplementary tables 1, 2 and 3. Enrichment analyses were conducted utilizing the fgsea package (v1.20.0) for Hallmark, YAP/TAZ and proliferation-associated markers gene sets through the fgseaMultilevel function, and the clusterProfiler (v4.14.6) package for GO gene sets via the gseGO function. Pathways below <0.05 p-adjusted were kept for visualization, and the results using org.Hs.eg.db v3.14.0 to map the gene name HGNC (Table S2). Finally, the outcomes were visualized using ggplot2 (v3.3.6) and plotted Enrichment plots via plot Enrichment from fgsea. To further analyze the enrichment pathways from GO, we utilized GOSemSim (v.2.32.0) to cluster related GO terms for classifying the overall group.(^91,92^)

### 4.8 Statistical analysis

In Figures 2 and 4, comparative studies were carried out by using the Kolmogorov-Smirnov test. Regarding experimental results shown in Figure 5, all data is expressed as means ± standard deviation (SD). Statistical analyses were performed using GraphPad Prism 10 software. The difference between groups was evaluated using ordinary one-way ANOVA followed by Tukey’s multiple comparisons test with a single pooled variance. In all figures, a p value lower than 0.05 was considered statistically significant with * p ≤ 0.05, **p ≤ 0.01 and ***p ≤ 0.001, **** p ≤ 0.0001. (ns), non-significative.

## Acknowledgments

Aldo Leal-Egaña and Christine Selhuber-Unkel acknowledge the financial support provided by the German Research Foundation (LE3418/4-1). Additionally, Aldo Leal-Egaña acknowledges the financial support provided by the FI EMS fund (Ex U. 6.1.22) and the 3DMM2O excellence cluter, Heidelberg University. Dr. Nicolás Tobar acknowledges the financial support provided by the Chilean National Research and Development Agency (ANID) grants: FONDECYT Initiation #11231225 and his ANID (Becas Chile #74220022). René Krüger was supported by the Federal Ministry of Education and Research (VIP+: PluripotencyScreen; 03VP11580). The authors thank the Microfabrication and Microfluidics Core Facility at the Institute for Molecular Systems Engineering and Advanced Materials (IMSEAM), which is funded partly by the Health + Life Science Alliance Heidelberg-Mannheim. All authors thank the strong support and participation of Andreas Vierling and Anastassia Fink in this research project.

---

**Supplementary Table 1.** Genes for cell dormancy.

| hgnc_symbol | Geneid | ENTREZID_id | Control_2D_1 | Control_2D_2 | Control_2D_3 | Released_cells_1 | Released_cells_2 | Released_cells_3 | Inside_capsules_1 | Inside_capsules_2 | Inside_capsules_3 | Source |
| --- | --- | --- | --- | --- | --- | --- | --- | --- | --- | --- | --- | --- |
| FHL2 | ENSG00000115641 | 2274 | 7.422537135 | 7.302601567 | 7.370765669 | 8.277169872 | 8.391822309 | 8.142911519 | 7.525193181 | 7.365465253 | 7.871397923 | https://www.science.org/doi/10.1126/sciadv.adr3997 |
| CTNNB1 | ENSG00000168036 | 1499 | 12.18138742 | 12.12296363 | 12.25410743 | 11.94361253 | 12.00424927 | 12.17432255 | 11.69820093 | 11.65473069 | 11.61065599 | https://www.tandfonline.com/doi/full/10.1080/28354311.2023.2264176#d1e593 |
| CBLB | ENSG00000114423 | 868 | 8.182516711 | 8.08448768 | 8.238906924 | 8.682365463 | 8.609981291 | 8.342950051 | 8.736058254 | 8.593917949 | 8.946553987 | https://www.tandfonline.com/doi/full/10.1080/28354311.2023.2264176#d1e592 |
| LIFR | ENSG00000113594 | 3977 | 7.186129904 | 7.151120289 | 7.572199595 | 8.02219511 | 8.310467522 | 7.980214619 | 7.78702442 | 7.992272227 | 7.690753548 | https://epub.uni-regensburg.de/40758/1/Ana%20Grujovic%20PhD%20thesis.pdf |
| CDKN1A | ENSG00000124762 | 1026 | 9.876364923 | 9.865807458 | 9.746359879 | 10.8737667 | 10.5916537 | 11.0893172 | 11.47924498 | 11.34312984 | 11.86960645 | https://www.science.org/doi/10.1126/sciadv.adr3997 |
| TNFAIP3 | ENSG00000118503 | 7128 | 7.173946324 | 7.072262547 | 7.183882601 | 7.872771728 | 7.789971197 | 8.0097824 | 7.372732584 | 7.355232823 | 7.864662338 | https://www.tandfonline.com/doi/full/10.1080/28354311.2023.2264176#d1e604 |
| EGFR | ENSG00000146648 | 1956 | 8.442511151 | 8.394549743 | 8.386964668 | 7.872771728 | 7.903617704 | 7.758154813 | 7.971848131 | 8.021770033 | 8.442040173 | https://www.tandfonline.com/doi/full/10.1080/28354311.2023.2264176#d1e606 |
| CDK6 | ENSG00000105810 | 1021 | 10.68570958 | 10.48334027 | 10.80090013 | 8.605151512 | 8.617425388 | 8.872243681 | 9.319810049 | 9.288015049 | 9.681557586 | https://www.tandfonline.com/doi/full/10.1080/28354311.2023.2264176#d1e614 |
| CUX1 | ENSG00000257923 | 1523 | 11.12427021 | 11.07295711 | 11.03102782 | 11.35140252 | 11.2842228 | 11.22716996 | 11.59404087 | 11.73581139 | 11.81415394 | https://www.tandfonline.com/doi/full/10.1080/28354311.2023.2264176#d1e594 |
| MYC | ENSG00000136997 | 4609 | 10.78395999 | 10.79216614 | 10.71521666 | 12.01390051 | 11.94862028 | 12.29811976 | 10.96196524 | 11.47645613 | 10.73932741 | https://www.tandfonline.com/doi/full/10.1080/28354311.2023.2264176#d1e600 |
| NDRG1 | ENSG00000104419 | 10397 | 8.90949568 | 8.60161439 | 8.637989197 | 10.85762501 | 11.18406499 | 10.36210704 | 9.876995223 | 9.727417539 | 10.06641733 | https://doi.org/10.1016/j.bbadis.2019.04.007 |
| COL17A1 | ENSG00000065618 | 1308 | 6.480352352 | 6.446638891 | 6.54142319 | 6.820986135 | 6.758683301 | 6.693327586 | 6.743114286 | 6.854671555 | 6.826759155 | https://link.springer.com/article/10.1186/s12943-025-02499-6 |
| H2AX | ENSG00000188486 | 3014 | 9.208288426 | 9.362100969 | 9.123859247 | 9.835856142 | 9.45982044 | 9.982638124 | 9.183824488 | 9.117086869 | 9.485152294 | https://link.springer.com/article/10.1186/s12943-025-02499-8 |
| PLK1 | ENSG00000166851 | 5347 | 9.716220496 | 9.644195128 | 9.697609669 | 9.871638751 | 9.597186932 | 10.00350763 | 9.435493644 | 9.207578833 | 9.37462218 | https://link.springer.com/article/10.1186/s12943-025-02499-13 |
| SOX9 | ENSG00000125398 | 6662 | 6.366559301 | 6.367217316 | 6.54142319 | 7.476896315 | 7.739921219 | 7.276429275 | 6.975820482 | 6.717716756 | 7.30481353 | https://link.springer.com/article/10.1186/s12943-025-02499-4 |
| PCNA | ENSG00000132646 | 5111 | 11.09019425 | 10.94879298 | 11.03237752 | 10.72152684 | 10.71088327 | 11.04183488 | 10.32253721 | 10.20617786 | 10.43299758 | https://link.springer.com/article/10.1186/s12943-025-02499-18 |

**Supplementary Table 2.**
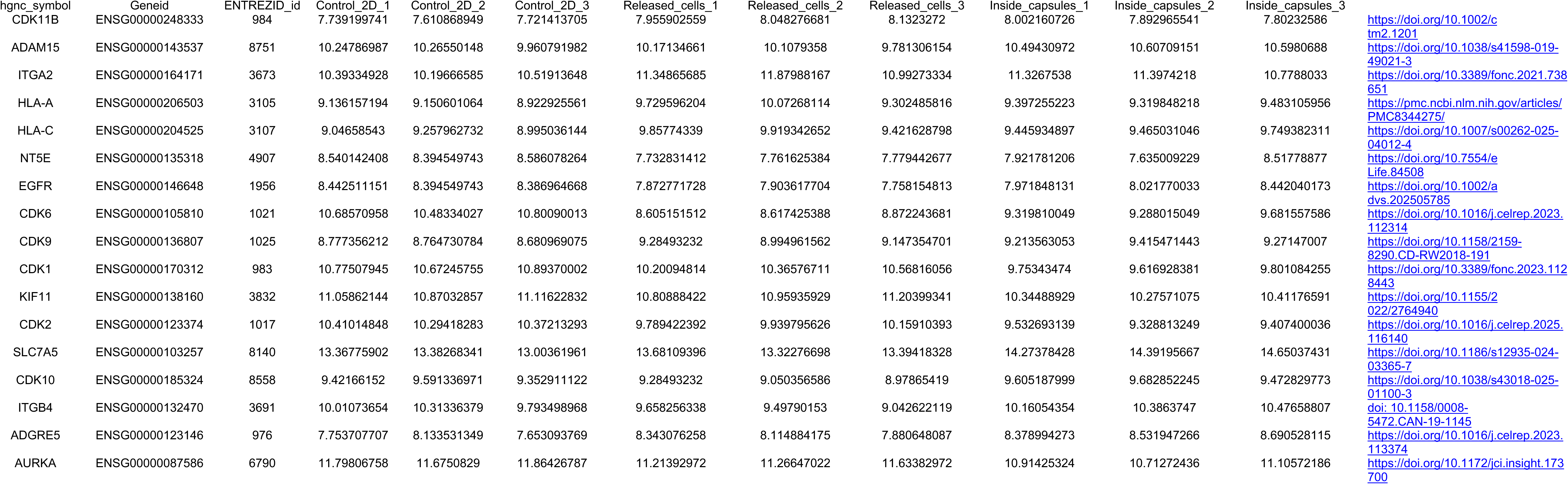
Genes for cell immunosurveillance.

**Supplementary Table 3.** Genes for cell early metastasis.

| hgnc_symbol | Geneid | ENTREZID_id | Control_2D_1 | Control_2D_2 | Control_2D_3 | Released_cells_1 | Released_cells_2 | Released_cells_3 | Inside_capsules_1 | Inside_capsules_2 | Inside_capsules_3 |  |
| --- | --- | --- | --- | --- | --- | --- | --- | --- | --- | --- | --- | --- |
| PIK3C2B | ENSG00000133056 | 5287 | 9.627055512 | 9.595046066 | 9.457908483 | 9.19384313 | 8.957740483 | 9.064203655 | 9.427084926 | 9.508988355 | 9.198262392 | https://doi.org/10.1016/j.celrep.2020.108105 |
| IKZF2 | ENSG00000030419 | 22807 | 9.306614158 | 9.241420191 | 9.389490149 | 8.329225966 | 8.48088734 | 8.287473272 | 8.356353242 | 8.450278458 | 8.597270994 | https://doi.org/10.1016/j.celrep.2020.108105 |
| IRS1 | ENSG00000169047 | 3667 | 10.82911949 | 10.84577543 | 10.82369274 | 12.30160537 | 12.52332559 | 12.11792145 | 11.82062569 | 12.05985331 | 11.94640398 | https://doi.org/10.1016/j.celrep.2020.108105 |
| ABCC5 | ENSG00000114770 | 10057 | 11.17667837 | 11.1780741 | 11.29831451 | 10.50656082 | 10.37114503 | 10.2537918 | 11.18616201 | 11.248029 | 11.56290627 | https://doi.org/10.1016/j.celrep.2020.108105 |
| CCNG2 | ENSG00000138764 | 901 | 8.300225193 | 8.317788513 | 8.451092794 | 9.223248865 | 9.252357562 | 8.660198605 | 8.920079915 | 9.096161034 | 9.228003148 | https://doi.org/10.1016/j.celrep.2020.108105 |
| ARHGAP10 | ENSG00000071205 | 79658 | 9.746404512 | 9.674292504 | 9.761156589 | 9.143426003 | 8.980762945 | 9.324907852 | 9.326616865 | 9.404924792 | 9.422440644 | https://doi.org/10.1038/nature20785 |
| CPE | ENSG00000109472 | 1363 | 8.425530177 | 8.411988589 | 8.301059122 | 9.645711241 | 10.24322338 | 9.342592905 | 9.135414248 | 8.997925116 | 9.002374291 | https://doi.org/10.1038/nature20785 |
| FAM13A | ENSG00000138640 | 10144 | 6.512419827 | 6.571893321 | 6.569749729 | 6.958225186 | 7.057035822 | 6.833785717 | 6.60218274 | 6.624589277 | 6.807598487 | https://doi.org/10.1016/j.celrep.2020.108105 |
| ADAM22 | ENSG00000008277 | 53616 | 8.743943854 | 8.646070996 | 8.729469126 | 8.242944881 | 8.409248079 | 8.28274382 | 8.246936199 | 8.539849766 | 8.246641307 | https://doi.org/10.1038/nature20785 |
| TRAM2 | ENSG00000065308 | 9697 | 10.1360987 | 10.08416159 | 10.11439137 | 9.394333459 | 9.401768979 | 9.446343346 | 9.439679376 | 9.450754197 | 9.562743485 | https://doi.org/10.1038/nature20785 |
| VEGFA | ENSG00000112715 | 7422 | 9.634254142 | 9.613447886 | 9.544284868 | 10.36459648 | 10.69468873 | 10.27794471 | 10.61133757 | 10.46209286 | 9.986684298 | https://doi.org/10.1016/j.celrep.2020.108105 |
| QRSL1 | ENSG00000130348 | 55278 | 9.19855636 | 9.278950666 | 9.387364405 | 8.859911724 | 8.858534173 | 9.13213434 | 8.803446867 | 8.692510296 | 8.982067642 | https://doi.org/10.1038/nature20785 |
| SYNE1 | ENSG00000131018 | 23345 | 7.332250939 | 7.415294929 | 7.319249923 | 8.062652455 | 8.059620206 | 7.793429823 | 7.902478675 | 8.23123767 | 7.634883354 | https://www.nature.com/articles/s44341-025-00018-6 |
| EGFR | ENSG00000146648 | 1956 | 8.442511151 | 8.394549743 | 8.386964668 | 7.872771728 | 7.903617704 | 7.758154813 | 7.971848131 | 8.021770033 | 8.442040173 | https://pmc.ncbi.nlm.nih.gov/articles/PMC4967137/ |
| VPS28 | ENSG00000160948 | 51160 | 9.060121989 | 9.14297393 | 8.79592157 | 9.259163823 | 9.121826124 | 9.090721463 | 9.512984701 | 9.631443045 | 9.32054569 | https://doi.org/10.3389/fmolb.2021.634183 |
| MLLT3 | ENSG00000171843 | 4300 | 8.257306816 | 8.221407312 | 8.386964668 | 8.543245863 | 8.748315978 | 8.756998102 | 8.309884626 | 8.416175171 | 8.361774868 | https://doi.org/10.1016/j.celrep.2020.108105 |
| P4HA1 | ENSG00000122884 | 5033 | 10.05018151 | 9.861204531 | 10.00835122 | 10.67752797 | 11.23854337 | 10.3545583 | 10.25094724 | 10.12417845 | 10.40426005 | https://doi.org/10.1016/j.celrep.2020.108105 |
| KLF4 | ENSG00000136826 | 9314 | 9.002367202 | 9.061635054 | 8.802394338 | 9.810424448 | 10.0405982 | 9.925450493 | 9.21845901 | 9.1300089 | 8.990806658 | https://doi.org/10.1038/s41420-023-01416-y |
| LCN2 | ENSG00000148346 | 3934 | 6.189314477 | 6.011397658 | 6.011397658 | 6.482870138 | 6.366724101 | 6.189002749 | 6.60218274 | 6.319719402 | 7.011221935 | https://doi.org/10.1038/nature20785 |
| MVB12B | ENSG00000196814 | 89853 | 8.04175073 | 8.175622078 | 7.779027605 | 7.574343227 | 7.413479841 | 7.597446892 | 7.516256124 | 7.712380526 | 7.917551595 | https://dx.doi.org/10.21037/tcr-2025-1498 |
| BAZ2A | ENSG00000076108 | 11176 | 10.10931616 | 10.12179895 | 9.99591502 | 10.72999846 | 10.58888855 | 10.55599514 | 10.7657968 | 10.85484344 | 10.76129886 | https://doi.org/10.1038/nature20785 |
| AHNAK | ENSG00000124942 | 79026 | 13.26611982 | 13.20216459 | 13.34988662 | 14.31159689 | 14.39831123 | 13.51214022 | 14.46906006 | 15.00682815 | 14.67247536 | https://doi.org/10.1038/nature20785 |
| CCDC15 | ENSG00000149548 | 80071 | 7.946760898 | 7.847020064 | 8.077600618 | 7.69516228 | 7.884306807 | 7.820931668 | 7.470467886 | 7.490329241 | 7.477431025 | https://doi.org/10.1038/s44341-025-00018-2 |
| DIAPH3 | ENSG00000139734 | 81624 | 8.471722403 | 8.420623787 | 8.593616014 | 8.379322503 | 8.315119921 | 8.532647148 | 7.921781206 | 8.196039626 | 8.040785289 | https://doi.org/10.1038/nature20785 |
| GPX2 | ENSG00000176153 | 2877 | 6.968838702 | 7.200255104 | 6.997165047 | 7.458229254 | 7.442228044 | 7.439288467 | 7.489009566 | 7.355232823 | 7.150111109 | https://doi.org/10.1038/nature20785 |
| UBALD1 | ENSG00000153443 | 124402 | 7.558169587 | 7.742079918 | 7.297909223 | 7.885927156 | 7.871264174 | 7.949939646 | 7.842705771 | 7.792068109 | 7.816454573 | https://doi.org/10.1016/j.celrep.2020.108105 |
| ANGPTL4 | ENSG00000167772 | 51129 | 6.716477951 | 7.030116091 | 6.869113667 | 7.624182421 | 7.725231056 | 7.06859604 | 6.911373712 | 7.112455879 | 7.271180049 | https://doi.org/10.1038/nature20785 |
| PLAUR | ENSG00000011422 | 5329 | 7.042545894 | 6.818196477 | 6.919532863 | 7.504320864 | 7.600657198 | 7.121736265 | 7.265074266 | 7.280654555 | 7.30481353 | https://pmc.ncbi.nlm.nih.gov/articles/PMC4967137/ |
| KCNG1 | ENSG00000026559 | 3755 | 7.412873252 | 7.62708718 | 7.095277987 | 7.061992807 | 6.969269505 | 7.208058133 | 7.064139098 | 7.13835671 | 6.881457634 | https://doi.org/10.1038/nature20785 |
| SPAG4 | ENSG00000061656 | 6676 | 7.412873252 | 7.472117421 | 7.242522213 | 7.949700063 | 8.059620206 | 7.589215835 | 7.801184311 | 7.770851447 | 7.974496329 | https://doi.org/10.1038/s44341-025-00018-2 |
| POFUT1 | ENSG00000101346 | 23509 | 10.07280914 | 9.974893912 | 10.0383036 | 9.519686959 | 9.710578176 | 9.464600649 | 9.765119986 | 9.862564506 | 9.889984968 | https://doi.org/10.1016/j.celrep.2020.108105 |
| CHMP4B | ENSG00000101421 | 128866 | 8.341798501 | 8.270476112 | 8.219178734 | 9.003182984 | 8.886299618 | 8.780171462 | 8.676067791 | 8.696029158 | 8.546138933 | https://doi.org/10.1038/s41392-022-01132-6 |
| APLP1 | ENSG00000105290 | 333 | 7.161606564 | 7.058433788 | 7.026326725 | 7.522237154 | 7.592412405 | 7.476416795 | 7.4227004 | 7.405438662 | 7.419618904 | https://doi.org/10.1038/nature20785 |
| TGFB1 | ENSG00000105329 | 7040 | 7.352988673 | 7.344987489 | 7.146921995 | 7.974329101 | 7.864691037 | 8.066936555 | 7.996155894 | 8.056289935 | 7.773567422 | https://pmc.ncbi.nlm.nih.gov/articles/PMC4967137/ |
| TNPO2 | ENSG00000105576 | 30000 | 10.35047082 | 10.3400137 | 10.22854267 | 10.53586059 | 10.32311326 | 10.27908477 | 10.92318146 | 10.944997 | 10.63149656 | https://doi.org/10.1038/nature20785 |
| SLC12A5 | ENSG00000124140 | 57468 | 7.909212583 | 8.067716705 | 7.886674148 | 7.548588809 | 7.550276632 | 7.605619683 | 7.794124943 | 7.968170163 | 7.576204113 | https://doi.org/10.1038/nature20785 |
| GDF15 | ENSG00000130513 | 9518 | 8.309576718 | 8.340827343 | 8.131851248 | 9.348479301 | 9.469941267 | 9.023461492 | 9.780004624 | 9.674119911 | 10.63149656 | https://doi.org/10.1016/j.celrep.2020.108105 |

**Supplementary Table 4.**
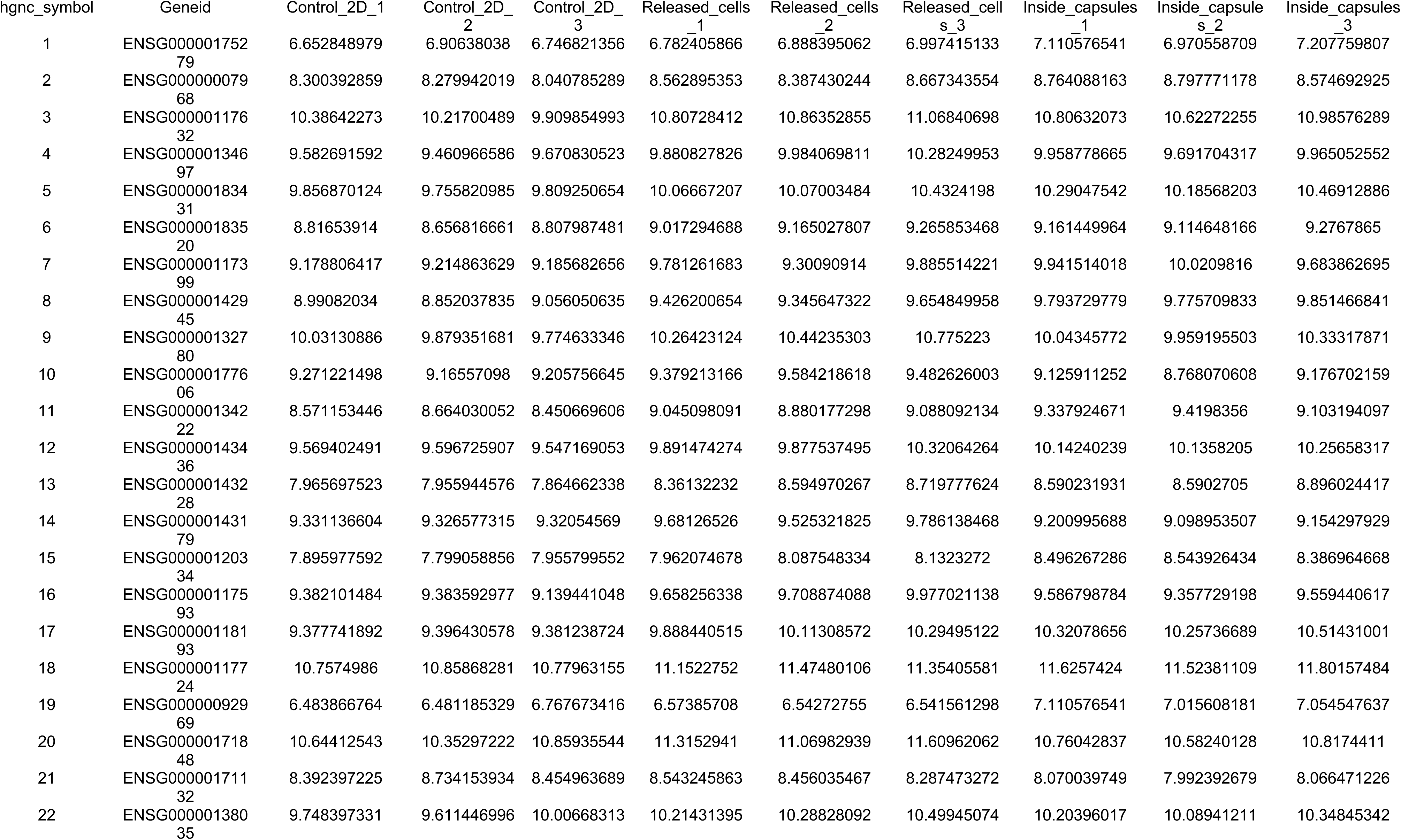
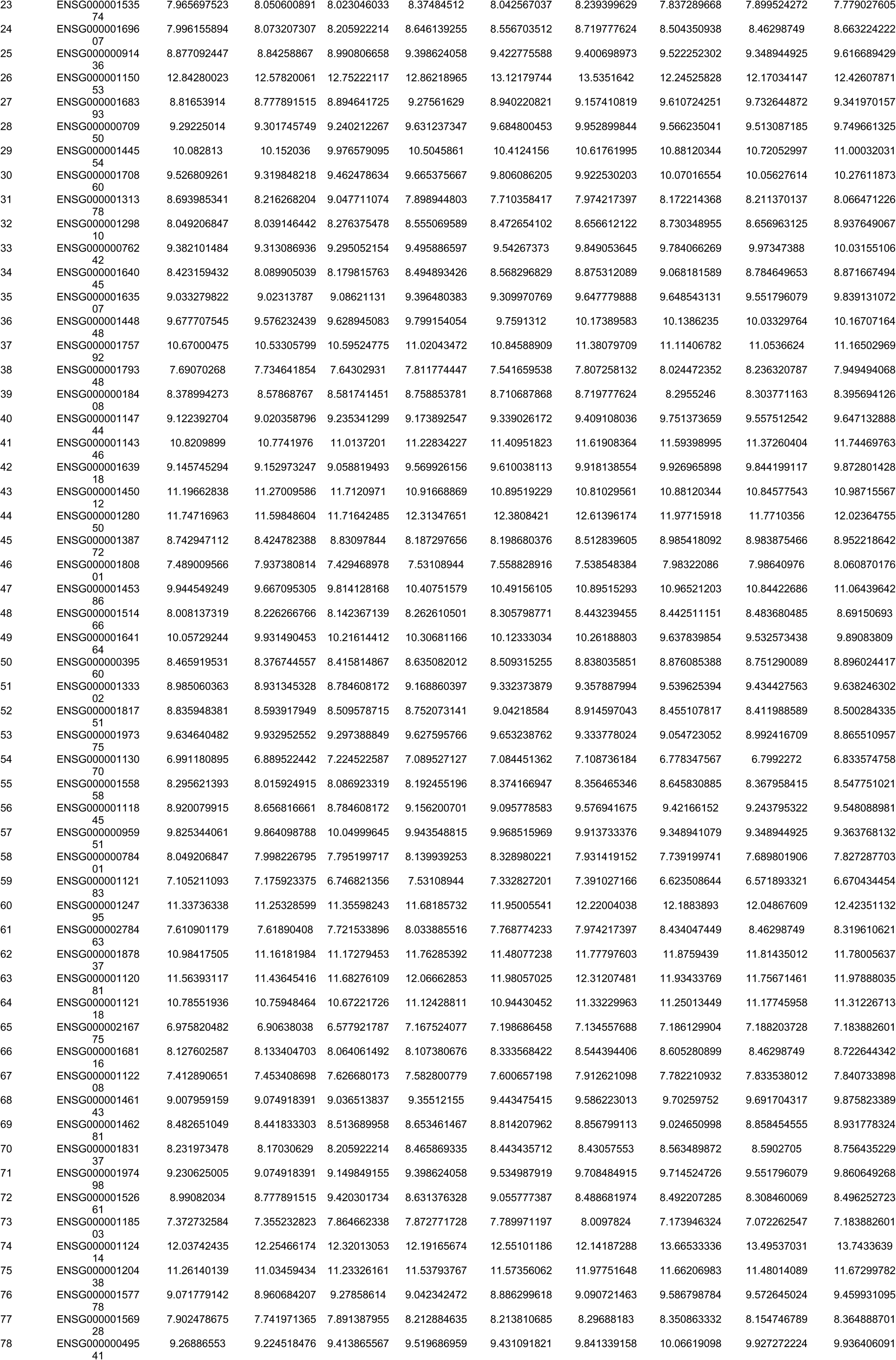
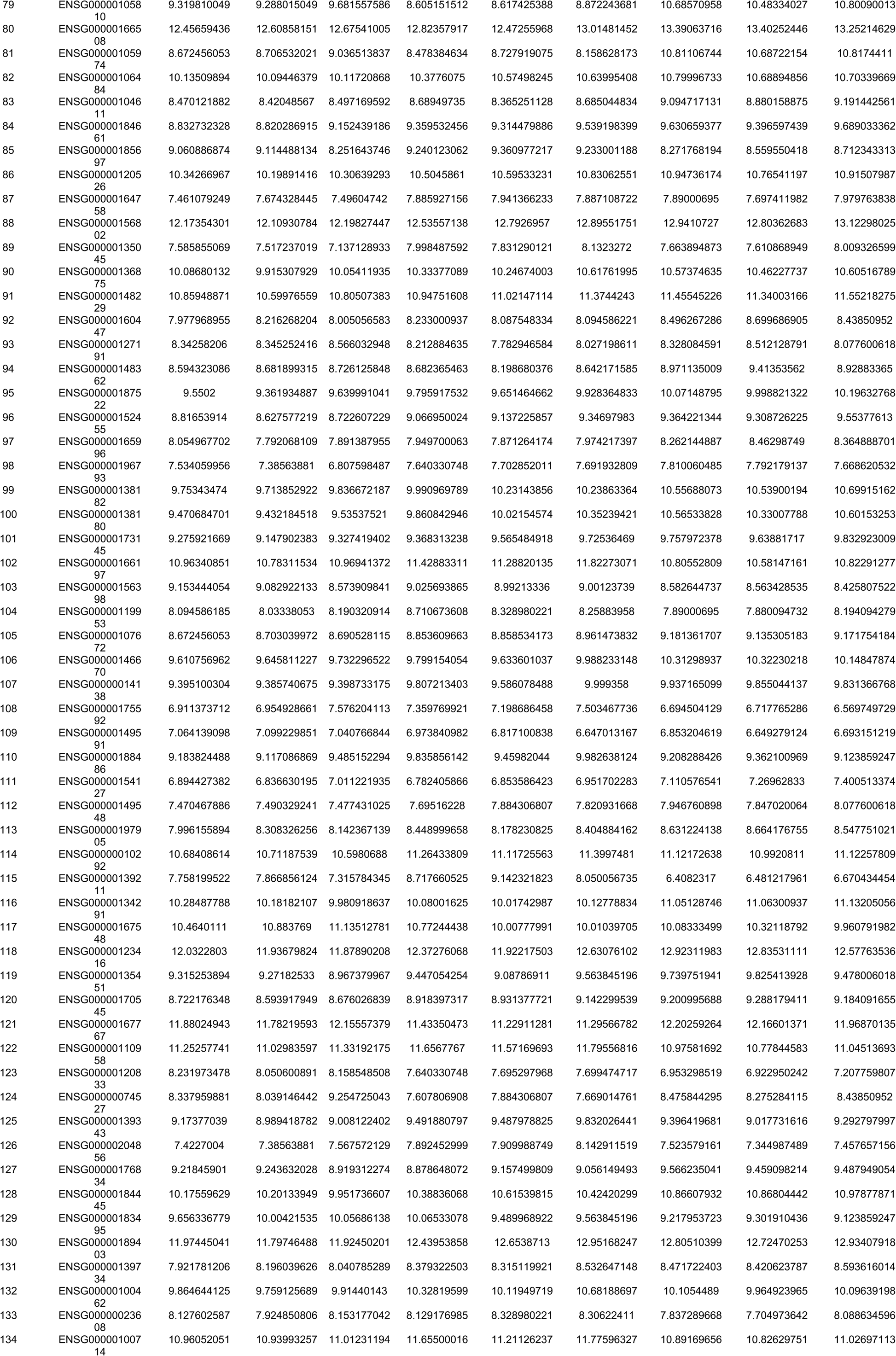
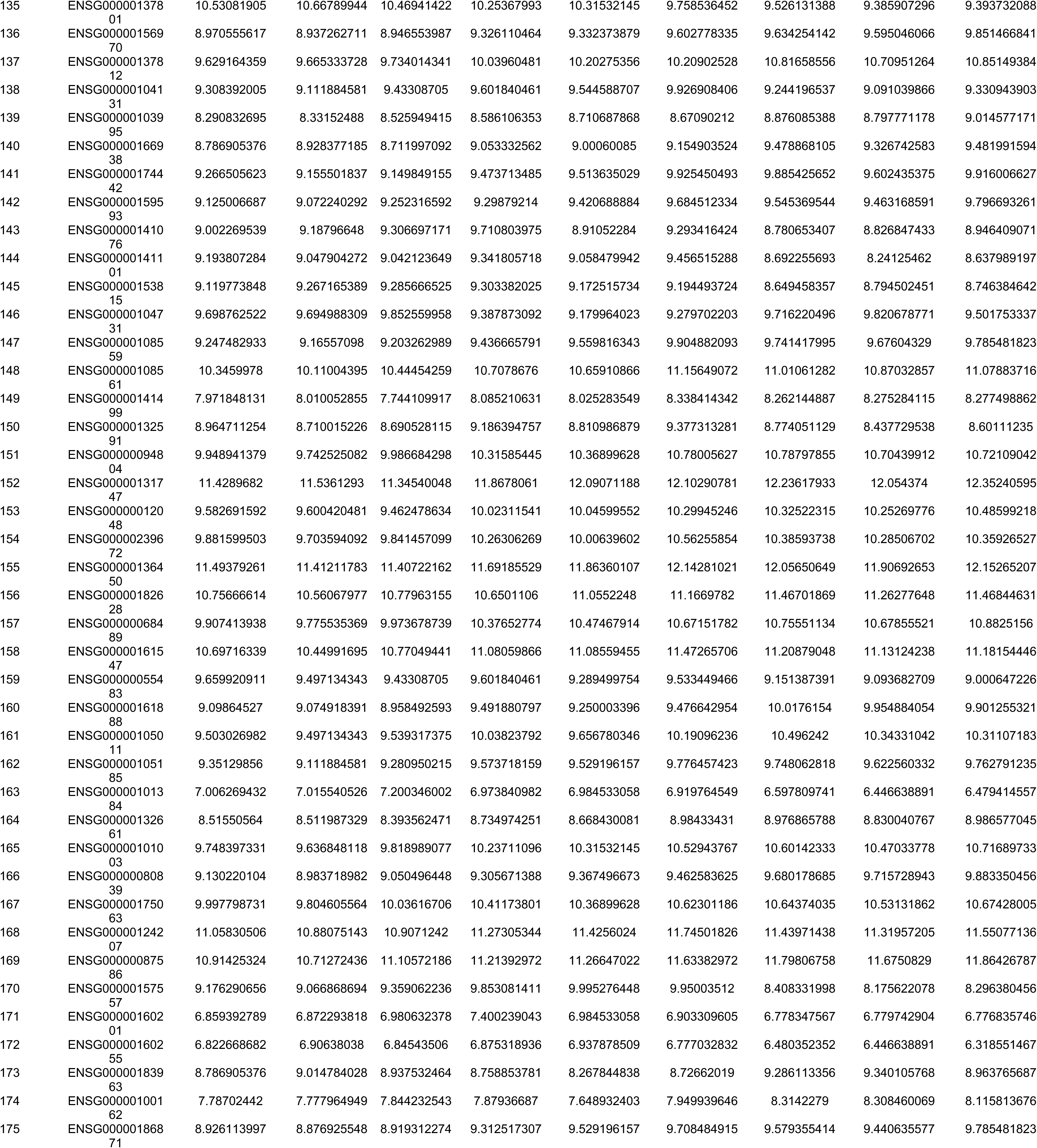
Yap / Taz Pathway.

**Supplementary Table 5.**
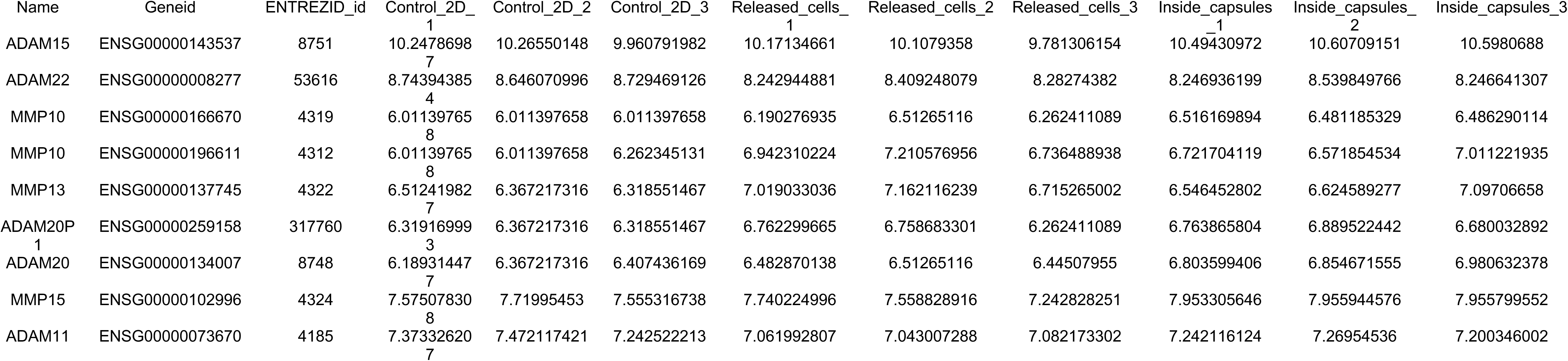
ADAMS and MMPs.

